# Open Blink: Low-cost TIRF microscopy for super-resolution imaging via *µ*Manager

**DOI:** 10.64898/2026.03.10.710894

**Authors:** Ran Huo, Jelle Komen, Moritz Engelhardt, Anaïs Millot, Jérôme Extermann, Kristin Grußmayer

## Abstract

Super-resolution localization microscopy (SMLM) has become a central tool for nanoscale biological research for its high spatial resolution and compatibility with wide-field microscopy. Achieving quantitative SMLM, however, requires homogeneous high-power illumination, nanometric axial stability, and precise multi-channel detection, features typically restricted to high-end commercial instruments or custom solutions in specialized laboratories. The cost of such microscopes and their technical complexity still limit the accessibility of these advanced imaging techniques. Several home-made single molecule microscopes and their submodules have been demonstrated as opensource, highly-customizable, and cost-effective alternatives for their commercial counterparts. Yet, implementation of such systems often requires expert knowledge in optics, electronics, and control system engineering. We introduce Open Blink, a compact open-source TIRF microscope integrating powerful homogeneous quad-line laser illumination, dual-channel detection, and active focus-lock stabilization for quantitative multi-color super-resolution imaging. Open Blink achieves a localization precision below 10 nm in dSTORM, supports a tunable, large field of view from 105 × 105 µm^2^ up to 257 × 257 µm^2^, and maintains axial stability over hours, enabling high-throughput super-resolution acquisition. Built with predominantly off-the-shelf components, and full integration into the open-source software *µ*Manager where metadata registration ensures reproducibility, Open Blink offers a low threshold for adoption by easing implementation, use and maintenance. At a substantially reduced cost of approximately 70 000 Euros, among which the high-power laser combiner alone is less than 20 000 euros, Open Blink greatly improves accessibility for laboratories who wish to implement scalable high performance super-resolution microscopy based on single molecules.

## 1 Introduction

Super-resolution microscopy techniques revealing information at nanometer scale have contributed to many new discoveries in life science.^1,2^ Among them, single-molecule localization microscopy (SMLM) and super-resolution optical fluctuation imaging (SOFI), based on the stochastic blinking of fluorophores, gained wide popularity because of their high resolution and compatibility with wide-field microscopes.^3–6^ Quantitative SMLM requires nanometric stability, homogeneous and high power illumination over large field of view and precise multi-channel registration, achievable only in high-end instruments. In particular, maintaining uniform blinking kinetics and localization precision over large fields of view remain technically demanding. However, costly equipment and maintenance, in addition to the difficulty of modifying commercial microscopes, hinder the accessibility of SMLM for researchers. With the continuous advancements of super-resolution microscopy by expert laboratories in recent years, its wider adoption called for affordable and easy-to-use hardware and corresponding control software for SMLM. In response, several opensource microscopes have been developed,^7^ which either focus on sub-modules such as the excitation source,^8,9^ microscope body,^10–12^ camera,^13^ focus-lock module,^14^ and control software,^15^ or provide a full-system solution as in K2 TIRF,^16^ miEye,^17^ and a 3D SMLM setup.^18^ Those open microscopy projects have demonstrated the benefits of home-made microscopes enabling different types of single molecule imaging, yet typically do not offer fully-integrated open control software, and provide limited laser power or field of view. Nevertheless, the trade-offs between desirable functionalities and cost, as well as between the technical complexity and user friendliness remain a challenge for scientists considering home-made microscopy setups. For example, customized setups are frequently built for specific use cases, with the design dictated by already available components for convenience, and therefore not straightforward to be transferred to another application or another laboratory. Likewise, home-made control electronics and research software are often not designed or optimized for re-usability, demanding expertise in both electronics and programming when implementing them.

To address these challenges, we designed Open Blink guided by the objective of achieving high-performance SMLM while ensuring accessibility, ease of use and low maintenance. We made design choices favoring long-term mechanical and optical stability and selected components commonly available and widely acknowledged by the community. Engineering expertise needed in building such as machining and soldering is minimized and easily outsourced. We prioritized software accessibility and scalability and aimed for hardware control and image acquisition schemes that are user-friendly and capable of metadata recording for downstream imaging analysis and data reproducibility. Despite the low cost, Open Blink is equipped with state-of-the-art features for SMLM, including homogeneous illumination, total internal reflection (TIR), and focus stabilization.

Compared to existing solutions, Open Blink demonstrates substantial improvements in several aspects. Firstly, without compromising the illumination qualities and homogeneity, the multimode high-power semiconductor lasers greatly reduce the price of laser sources and offer a high-power open-hardware laser combiner alternative, as demonstrated in Section 2.1. Secondly, the homemade microscope body adapted from the open microscope frame miCube is economical, mechanically stable, and easy to modify.^11^ We further reduced the miCube frame size to enable an enlarged field of view (FOV) sized over 200 × 200 µm^2^, as shown in Section 2.2, while preserving illumination homogeneity and thus localization precision. Together with the focus-lock module ‘*fg*Focus’, which we developed based on *pg*Focus and *qg*Focus^16,19^ to address the absence of an up-to-date and *µ*Manager-compatible implementation, this makes Open Blink particularly suitable for longterm high-throughput single molecule based super-resolution microscopy powerful for biological discovery.

Most open hardware projects develop their customized control software with Python, Lab-VIEW, or C#.^9,16,17^ Instead of specialized programming and varying graphic interfaces, we opted for the open-source microscopy control software *µ*Manager for Open Blink, which is well-known to the bioimaging community for its integrated functions for common microscope hardware control and imaging acquisition tasks. A large number of device adapters from hardware vendors and developers, and customizable development capability enabled by pycroManager, pymmcore-plus and EMU^20–22^ make *µ*Manager flexible for extension with minimal specialized programming, which we demonstrate in Section 2.3 with our laser combiner controller and focus-lock controller *µ*Manager plugins. Communication between hardware and software for Open Blink is established through simple micro-controllers without complex electronics. Metadata containing imaging parameters such as stage positions, laser power and exposure time are also incorporated for each image acquired, ensuring appropriate documentation for reliable image analysis and reproducible experiments.

Finally, we showcase applications of Open Blink in single molecule super-resolution cell imaging with DNA-PAINT, dSTORM and SOFI respectively in Section 2.4.

In total, Open Blink cut the cost of a full super-resolution microscope system significantly to approximately 70 000 Euros. Documentation of Open Blink, including the list of components, design files, and compiled ready-to-use *µ*Manager plugins, is available at Grußmayer Lab GitHub page. With semi-turnkey components, simple control electronics, and full *µ*Manager integration, Open Blink provides a reproducible and scalable platform for high-performance quantitative single-molecule super-resolution microscopy at substantially reduced cost.

## 2 Results

### 2.1 Low-cost high-power laser combiner with homogeneous illumination

Single-molecule localization microscopy (SMLM) reconstructs a super-resolved image by precisely localizing the position of individual molecules in tens to hundreds of thousands of images.^23–25^ Implementations such as direct stochastic optical reconstruction microscopy (dSTORM) demand high power laser excitation up to 10 kW/cm^2^ for photoswitching of organic dyes to achieve sparse single-molecule conditions.^26,27^ High power excitation sources from single mode lasers comprise a large cost of SMLM setups. Inspired by Ries lab’s laser engine,^8^ which employed high power-cost ratio laser diodes with customized drivers, we built a quad-line laser combiner with semi-turnkey diode lasers. In a similar layout as Ries lab laser engine, two diode lasers at 638 nm are combined via a polarizing beam splitter to double the excitation power at the far-red wavelength, in parallel to the 405 nm and 488 nm diode lasers (LAB-405-500/LAB-488-2000/LAB-638-1000, Lasertack GmBH), as shown in Fig.1 (a) and Fig.S1 in the supplementary information. A diode-pumped solid-state (DPSS) laser at 561 nm (MGL-FN-561, CNI Laser) completes the visible spectrum coverage. All combined laser beams are launched into a multimode optical fiber (MMF) with an aspherical lens.

**Fig 1.**
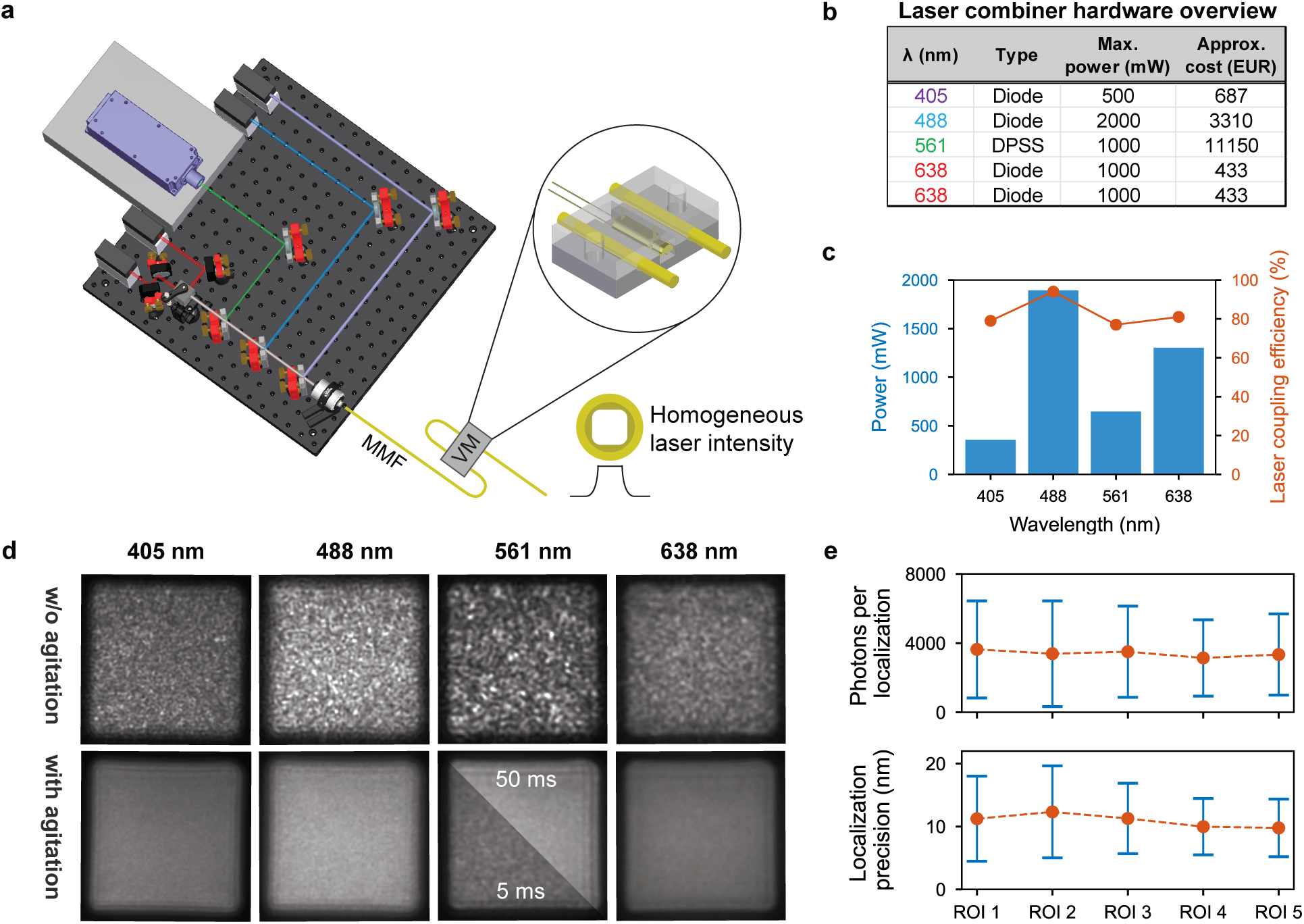
A low-cost high-power laser combiner with multimode fiber (MMF) pigtail delivers homogeneous illumination. **(a)** 3D representation of the quad-line laser combiner of Open Blink; agitation from vibrational motor (VM) is transmitted to the MMF in a customized mount displayed in the zoom-in image. **(b)** List of the lasers used, their power, and approximate prices. DPSS: Diode-pumped solid-state laser. **(c)** Power performance of the laser combiner at four wavelengths, shown as laser coupling efficiency (orange line) and maximum output power (blue bar). **(d)** Comparison of laser intensity profile at the MMF output in the presence and absence of agitation from the VM. Images taken directly at the MMF exit with 5 ms exposure time except for the insert image for 561 nm which has exposure time of 50 ms. **(e)** Average photon count per localization (top) and localization precision (bottom) of DNA-PAINT imaging at five different regions of interests (ROIs) across the illuminated area at the sample plane. Error bars indicate the standard deviation from all localizations in each ROI.

Usage of the semi-turnkey diode lasers is much more cost-effective compared to single mode lasers of similar power (summarized in Fig.1 (b)), amounting to ∼29% of the total cost of Open Blink. When compared to using unmounted and uncollimated laser diodes, it also saves one from soldering work and laser driver current calibration. Although the diode lasers selected are supplied with calibrated drivers, they lack modulation units or control software. We developed the control electronics and *µ*Manager plugin for the Open Blink laser combiner based on micro-controllers which is detailed in Section 2.3.

SMLM image quality is substantially affected by the homogeneity of excitation across the field of view, as the localization precision is proportional to the number of detected photons, which in turn depends on the excitation laser intensity.^28^ The laser excitation also influences the photoswitching or photoactivation behavior of fluorophores, and inhomogeneities in the illumination can lead to variations in molecular densities. A homogeneous illumination thus ensures a consistent resolution over the field of view, crucial for interpreting SMLM rendered images.^27^ A typical MMF has an inhomogeneous output intensity profile patterned with speckles due to the interference of all propagating modes. It has been reported that a mode scrambler effectively reduces such speckles, thus achieving homogeneous illumination at the MMF output.^8–10^ In a similar design reported by Lam et al.,^9^ we used a customized MMF (NA 0.22, CeramOptec), whose square core profile of size 70 µm by 70 µm enables TIRF illumination (detailed in Section 2.2) and maximizes the effective detection area on the camera chip. The homogeneity of laser illumination is facilitated by a vibrational motor (VM) agitating the MMF connected through a 3D-printed mount (polylactic acid, CAD available), as shown in the zoom-in schematics in Fig.1 (a).

Compared to single mode optical fibers, MMFs have higher numerical apertures (NA), which eases the coupling of input lasers. We measured the MMF coupling efficiency and the maximum output power of the laser combiner at the exit of the MMF. The coupling efficiency for different wavelengths varies between 77% to 94% (Fig.1 (c)), calculated by the ratio of laser intensity exiting and entering the MMF. These values demonstrate that high-power multimode excitation can be achieved with efficiencies comparable to single-mode fiber delivery while enabling substantially reduced system cost. Notably, at 638 nm the laser combiner yields a maximum power of 1.3 W or intensity of 6.94 kW/cm^2^, sufficient for high-quality SMLM (example image shown in Section 2.4). The homogeneity of the intensity is first verified by imaging the MMF output intensity profile (Fig.1 (d)). At 5 ms exposure time, the diode lasers at 408 nm, 488 nm, and 638 nm exhibit significantly decreased speckles. The more coherent single mode DPSS laser at 561 nm results in more speckles at this fast image acquisition rate, which can be effectively reduced at increased exposure times of 50 ms, well within the typical range of exposure times used for single molecule imaging. Note that, among other MMFs we have tested, the performance of homogeneity from mode scrambling by vibrational motor varies. A small core MMF (M42L05, Thorlabs) compatible with TIRF as used in Kwakwa et al.^10^ does not respond to the agitation, while a larger square core MMF (M103L05, Thorlabs) as used in Schroder et al.^8^ shows improved homogeneity when agitated. Images of their respective output intensity profile are found in Fig.S2 of the supplementary information. We suspect that the variations in the coatings applied around the cladding of the MMFs affect the effectiveness of mode scrambling. The more rigid polymer coating in the Thorlabs M103L05 might subject the MMF to increased microbends compared to the acrylate coatings used in the M42L05, thus inducing effective spatial mixing of the propagation modes.^29^

Next, we examined the effect of homogeneous illumination for super-resolution imaging quality by DNA-PAINT imaging (details in Section 2.4). The sample was imaged under the same laser intensity at five different regions of interest (ROIs) sized 15.6 *µ*m *×* 15.6 *µ*m distributed across the full illuminated area. The positions of the ROIs are indicated in the Fig.S3 in the supplementary information. Fig.1 (e) shows the average photon count per localization and the localization precision which reflect the excitation intensity in each ROI. The spatial consistency of photon counts and localization precision across all ROIs demonstrates homogeneous excitation over the full illuminated field of view, ensuring spatially invariant resolution performance.

### 2.2 Home-built TIRF microscope body with large FOV and focus-lock module

MiCube^11^ is an open-source microscope body frame which we adopted for its mechanical stability, accessibility and customizability in comparison to commercial microscopes. It is compatible with brightfield imaging, widefield epifluorescence, highly inclined and laminated optical sheet (HILO) and total internal reflection fluorescence (TIRF) microscopy.^16^ TIRF is an effective optical sectioning method typically used to illuminate molecules immobilized on the coverslip or to visualize cellular structures on or in the vicinity of cell membranes at a high signal-to-background ratio (SBR). A small focused excitation beam size at the back focal plane (BFP) of a high-NA objective is critical to enabling TIRF. In practice, the TIRF condition is typically achieved by using a single mode laser with small beam diameter, or by laser delivery through single mode optical fibers. Lam et al.^9^ showed that instead of single mode lasers or single mode fibers, a collimated beam from a MMF with small core sized 70 *µ*m *×* 70 *µ*m can be focused by a focusing lens (FL) of 125 mm focal length to meet the TIRF condition.

On the other hand, a large field of view (FOV) in microscopic imaging is highly favorable for high-throughput imaging and studying cell-cell interactions when multiple targets need to be visualized simultaneously. To achieve this, the illumination area in the sample plane should be increased, which can be realized by enlarging the excitation beam size. However, the beam size is restricted by both the TIRF condition and the physical size of the optical components. Our design, combining the 70 *µ*m *×* 70 *µ*m core size MMF and a 150 mm focusing lens, results in a large illumination area of 257 *µ*m *×* 257 *µ*m, while preserving TIRF condition which is shown by detailed calculations in the supplementary information 4. At the same laser power, exposure time, and display contrast, we observed higher SBR for the same cell imaged in TIRF than in epi-illumination (Fig.2 (b)). To gain the flexibility to choose between a large FOV size and a high illumination intensity, a beam size reducer consisting of two achromatic lenses in a telescope configuration that halves the beam diameter can be added after the collimation lens at the MMF output (see Section 4). With the simple insertion or removal of the beam size reducer, the FOV can be switched between the small and large configuration easily. The halved beam size translates to a quartered FOV size and a four times higher power density at the sample plane, facilitating dSTORM and other power-demanding wide-field imaging. Table S1 in the supplementary information shows the laser intensity measured at the sample plane at the large and small FOV sizes.

**Fig 2.**
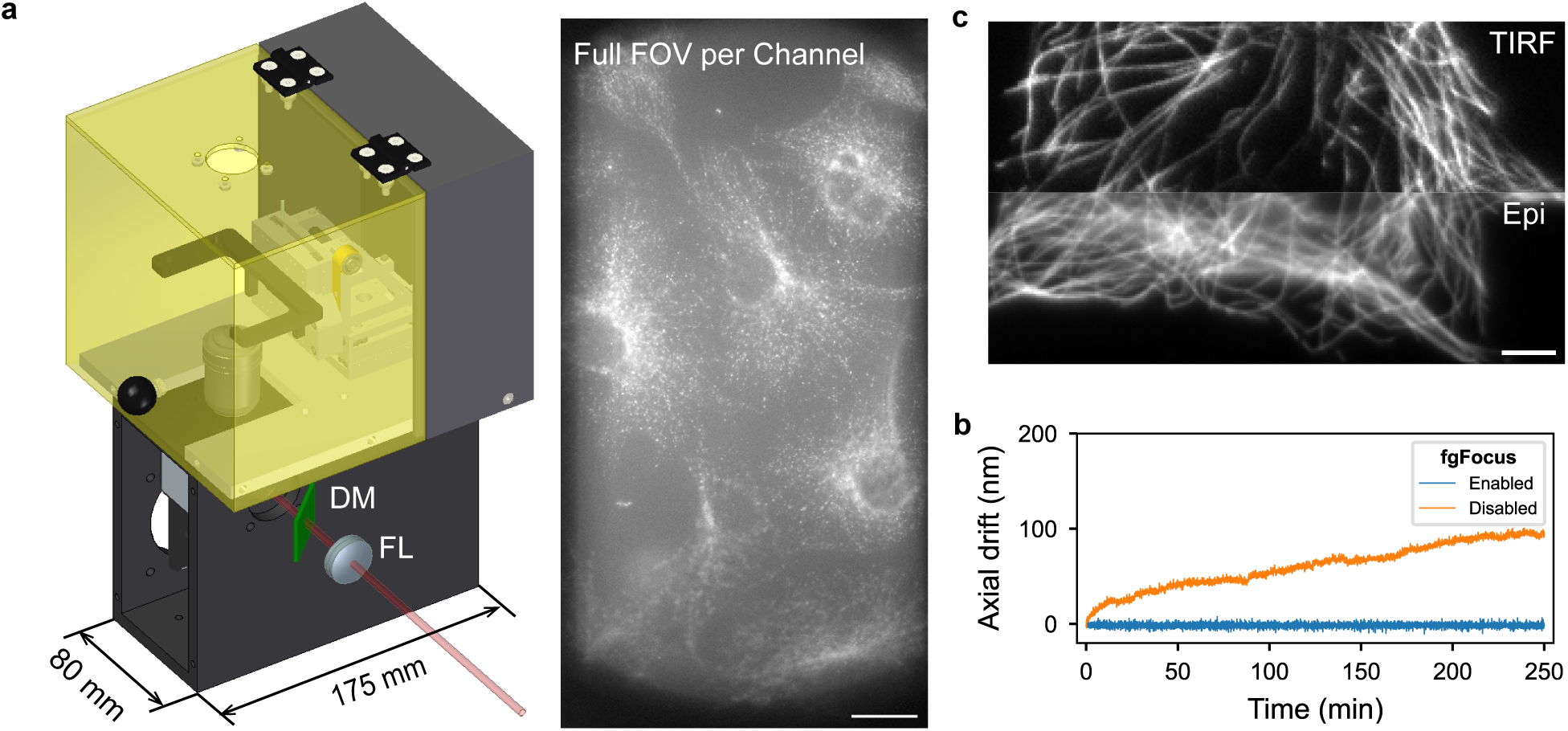
A compact microscope body with large FOV and focus-lock module. **(a)** Left: CAD representation of the modified MiCube with reduced width, accommodating 150 mm focusing lens (FL) and infrared dichroic mirror (DM). Right: the modification enables large FOV visualizing multiple cells in either of the dual-color detection. Scale bar = 20 *µ*m. **(b)** Comparison of cell images under TIRF (upper) and epi-illumination (lower). Scale bar = 5 *µ*m. Images were taken with identical laser power and exposure time, and displayed in the same contrast. **(c)** Comparison of axial drift with and without focus-lock module *fg*Focus.

The original miCube employs a 200 mm focusing lens that focuses the excitation beam onto the back focal plane. To accommodate the 150 mm focusing lens in our design, we reduced the width of the miCube to 80 mm (Fig.2 (a), left) in order to place the focusing lens closer to the miCube. Switching of the illumination modalities between epi-, HILO and TIRF is conveniently enabled by a motorized linear stage on top of which the focusing lens is mounted. The modified miCube accommodates a fixed 60*×* high-NA objective (UPLAPO60XOHR, Olympus) at the top. A compact sample-positioning stage (three CLS5252 assembled, SmarAct) providing nanometer precision and 31 mm travel range in each axis, is also mounted on top of miCube with an adapter plate in between (customized cut from aluminum, CAD available). The adapter plate extends above the miCube’s top surface and functions as the base for the enclosure needed for environmental control (customized cut acrylic plate, CAD available). The sample stage is compatible with a standard circular sample (25 mm diameter) or rectangular sample (35 mm *×* 76 mm) by attaching corresponding sample holder adapters to the stage.

Multicolor imaging expands the palette of usable fluorophores and enables simultaneous visualization of different molecules which is essential for studying complex cellular structures, protein interaction and cellular dynamics. The detection path of Open Blink is configured as dual-channel split by a dichroic mirror at 595 nm (ET595/50, Chroma). The two channels are imaged side by side on an sCMOS camera (BSI Express, Photometrics) after a 4f relay optics. Final magnification measures at 59.9 *×*, corresponding to a camera pixel size of 108.4 nm. An adjustable aperture is placed at the focal point of the tube lens (TTL180-A, Thorlabs) for restricting the image size per channel to half of the camera chip size. Fig.2 (a) right panel shows a snapshot of multiple cells imaged in one color channel under the large FOV configuration. The full schematic layout of the microscope can be found in Fig.S1 in supplementary information.

Super-resolution imaging based on single molecule blinking often requires tens of thousands of frames, leading to long acquisition times, during which focus drift may compromise the imaging quality. An active focus-lock system is beneficial for long time-lapse and automated superresolution imaging. We implemented a focus-lock module *fg*Focus, short for ‘fairly good’ and adapted from *pg*Focus and *qg*Focus.^16,19^ Maintaining of the focus is realized by coupling an infrared beam using a dichroic mirror to the microscope excitation path in TIRF configuration. The reflected infrared beam can be detected by a one-dimensional sensor array (TSL1401-DB, Parallex) and is used to determine the sample z-position. The shift of the infrared beam position on the sensor induced by sample drift is fed into a proportional–integral–derivative (PID) controller, which actuates the sample z-stage to compensate for the drift. In our design, the 150 mm focal length of the focusing lens for TIRF poses a challenge to fit in the dichroic mirror in between the focusing lens and the MiCube, since the original MiCube has a focusing lens at 200 mm for TIRF. We thus used a compact kinematic mirror mount (KMS/M, Thorlabs) onto which the dichroic mirror is fixed with silicone glue. *fg*Focus is made compatible with *µ*Manager and detailed in Section 2.3. The stability of *fg*Focus is demonstrated by imaging a fluorescent bead slide (T14792, Thermo Fisher) over 250 minutes. The drift of the focus reported by the infrared detector is traced every 2 seconds either with and without the activation of *fg*Focus. While enabling *fg*Focus, no drifting of focus was visually observable during the experiments, with the axial drift averages at -1.58 nm (*±* 2.00 nm standard deviation), significantly improved compared to the non-enabled case where the final axial drift accumulated to 94.2 nm, demonstrating nanometric axial stability over multi-hour acquisitions required for quantitative SMLM.

### 2.3 µManager integration

*µ*Manager is an open-source microscope control software integrating core microscopy functionalities such as stage control, camera settings and image acquisition tasks in its default graphic user interface (GUI). *µ*Manager supports a wide range of hardware components through the so-called device adapters that bridge hardware and software. The ease of use and flexibility of this modular approach to adapt customized microscopes motivated us to integrate full control and image acquisition for Open Blink in *µ*Manager. In addition, *µ*Manager is available under open source license for all common operating systems (Windows, Mac and Linux) and based on the widely used image processing environment ImageJ, which is particularly popular in the bioimaging community that we target as end users of Open Blink.

Although most components used in Open Blink from mainstream manufacturers come with device adapters for *µ*Manager, which include the sample stage, TIRF stage and sCMOS camera, the customized parts, i.e. the laser combiner, vibrational motor, and the infrared detection and z-stage feedback for *fg*Focus at Open Blink require tailored solutions taking care of both the control electronics and communication with *µ*Manager. The individual device demands different types of control signals. The speed of the vibrational motor can be controlled by pulse width modulation (PWM) signals. The intensity of all lasers is modulated by 0-5 V analog voltages. *fg*Focus, which relies on analysis of the beam profile detected by the infrared detector, needs to receive an analog readout from the infrared sensor. To achieve this, we employed two micro-controllers, which output or read appropriate signals to or from the devices through serial communication with *µ*Manager (Fig.3 (a)). Micro-controller 1 is the controller for the laser combiner. A customized PCB integrates a Raspberry Pi Pico, three 12-bit dual-channel digitalanalog-converters (DACs) and a MOSFET transistor. The DACs convert the digital signal from the micro-controller into analog voltages to the laser drivers, and the MOSFET modulated by the PWM signal is wired to the vibration motor, facilitating homogeneous illumination. Micro-controller 2 is a thumb-sized lower-power board (XIAO SAMD21, Mouser Electronics) which sends to *µ*Manager the analog signal read from the infrared sensor board (TSL1401-DB, Parallax). Both micro-controllers are connected to the computer via USB connection. The PCB design and configuration of both boards are available at https://github.com/GrussmayerLab/LaserController and https://github.com/GrussmayerLab/fgFocus, respectively.

**Fig 3.**
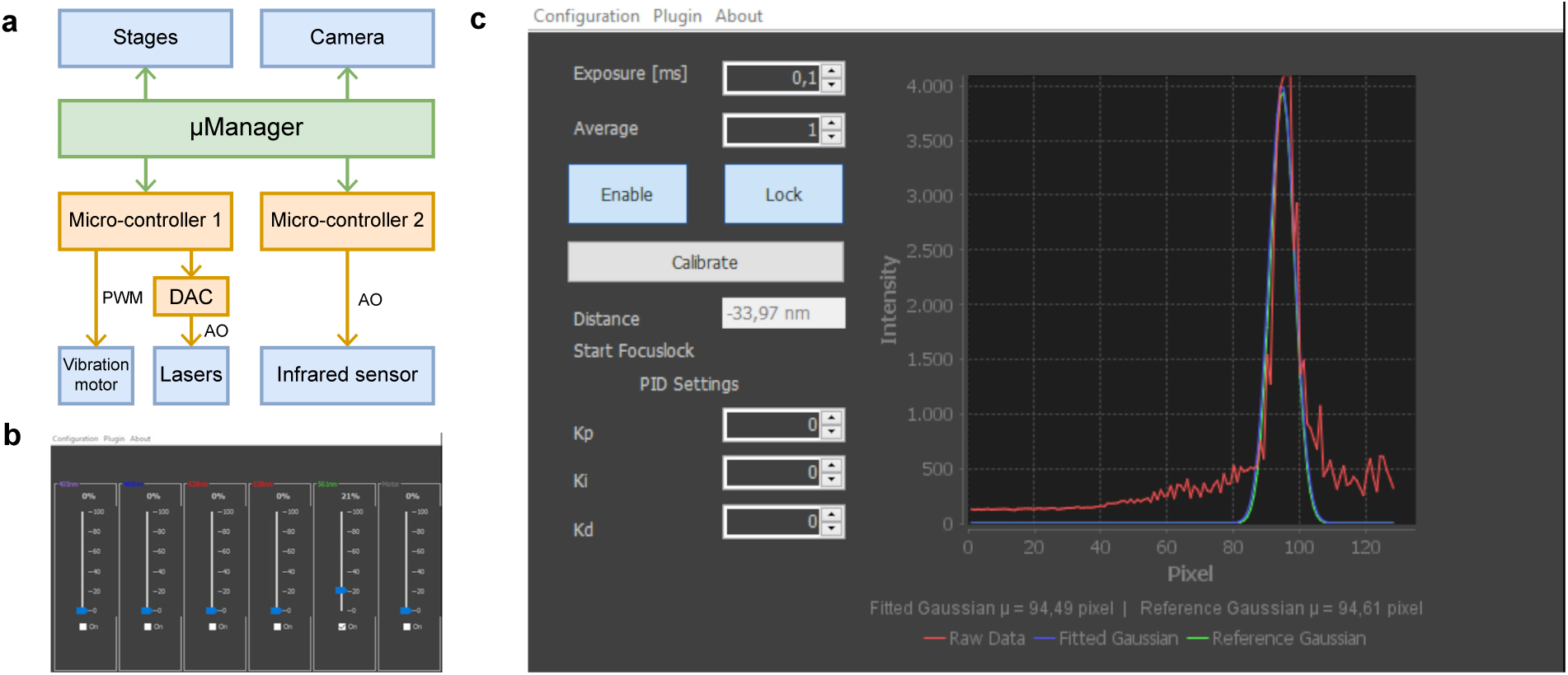
Open Blink integration in *µ*Manager. **(a)** Control scheme for customized components. Micro-controller 1 controls the laser intensity and vibrational motor speed of the laser combiner; micro-controller 2 reads the pixel values from the infrared sensor as the input for the PID control logic of the focus-lock module; **(b)** Laser controller plugin interface in *µ*Manager. Laser power and vibrational motor speed can be adjusted by the sliders; **(c)** *fg*Focus plugin interface in *µ*Manager which displays the infrared beam profile. Infrared sensor and PID controller parameters can be configured.

For software integration, existing device adapters of micro-controllers for *µ*Manager do not support general DAC output.^30^ Commercial data acquisition cards compatible with *µ*manager are pricey and vastly exceed the requirements for simple analog signal generation. They are also missing the essential ability of metadata writing, which is an advantage of integrating all instrumental control in one place in order to record the relevant device parameters in parallel with the image acquired in *µ*Manager. For the laser and vibrational motor controller, we bypass the typical hardware configuration in *µ*Manager, and instead use a plugin within the Easier Micro-manager User interface (EMU)^22^ facilitating metadata writing in a straightforward GUI (Fig.3 (b)). We provide a similar solution for *fg*Focus software which is the first *µ*Manager compatible focus-lock module for the increasingly popular SmarAct sample stage. Through an EMU plugin, shown in Fig.3 (c), the *fg*Focus GUI displays the reflected beam profile which is fitted with a Gaussion function. When in focus, *Calibrate* function scans *±*1 *µ*m from the current z-stage position to determine the linear relationship between the beam position on the sensor and the actual sample focal position. Once *Lock* is enabled, the current beam position on the sensor is used as the reference and *fg*Focus maintains the focus through proportional–integral–derivative (PID) control logic whose parameters can be configured within the plugin. Both EMU plugins for the laser combiner controller and *fg*Focus are pre-compiled and downloadable for immediate use in *µ*Manager in the aforementioned repositories.

### 2.4 Application in single molecule super-resolution imaging

Super-resolution imaging based on localizing single molecules relies on the sparse blinking condition of fluorophores, often enabled by high-intensity illumination as in dSTORM.^27^ We showcase the compatibility and versatility of Open Blink with three types of single molecule super-resolution imaging applications that have different illumination requirements: dSTORM, DNA-PAINT and SOFI for 2D high-resolution imaging of microtubules in fixed cells. Typical dSTORM fluorophores such as Alexa Fluor 647 require high laser power in combination with an oxygen scavenger and thiol buffer system to induce blinking,^25,26^ i.e. on- and off- switching of fluorescence. Therefore, we used the small FOV configuration in the excitation path at 1.2 kW/cm^2^, resulting in sparse single-molecule imaging and reached a localization precision of 8.21 nm, consistent with the expected photon-limited precision under high-power excitation (Fig.3 (a)). DNA-PAINT uses the reversible hybridization of DNA oligos, namely the docking strand coupled to the imaging target and the imager strand that is supplied in solution at low concentration, to achieve single-molecule binding events.^31^ It does not require light to induce blinking, but suffers from background noise arising from the imagers carrying fluorophores floating around the sample, and thus necessitates TIRF illumination for optimal performance. Fig.4 (b) shows a DNA-PAINT image using Cy3B imagers matching the P1 docking strand under the TIRF illumination and the large FOV configuration, with localization precision estimated at 13.0 nm, despite operating under large-FOV TIRF illumination conditions.

**Fig 4.**
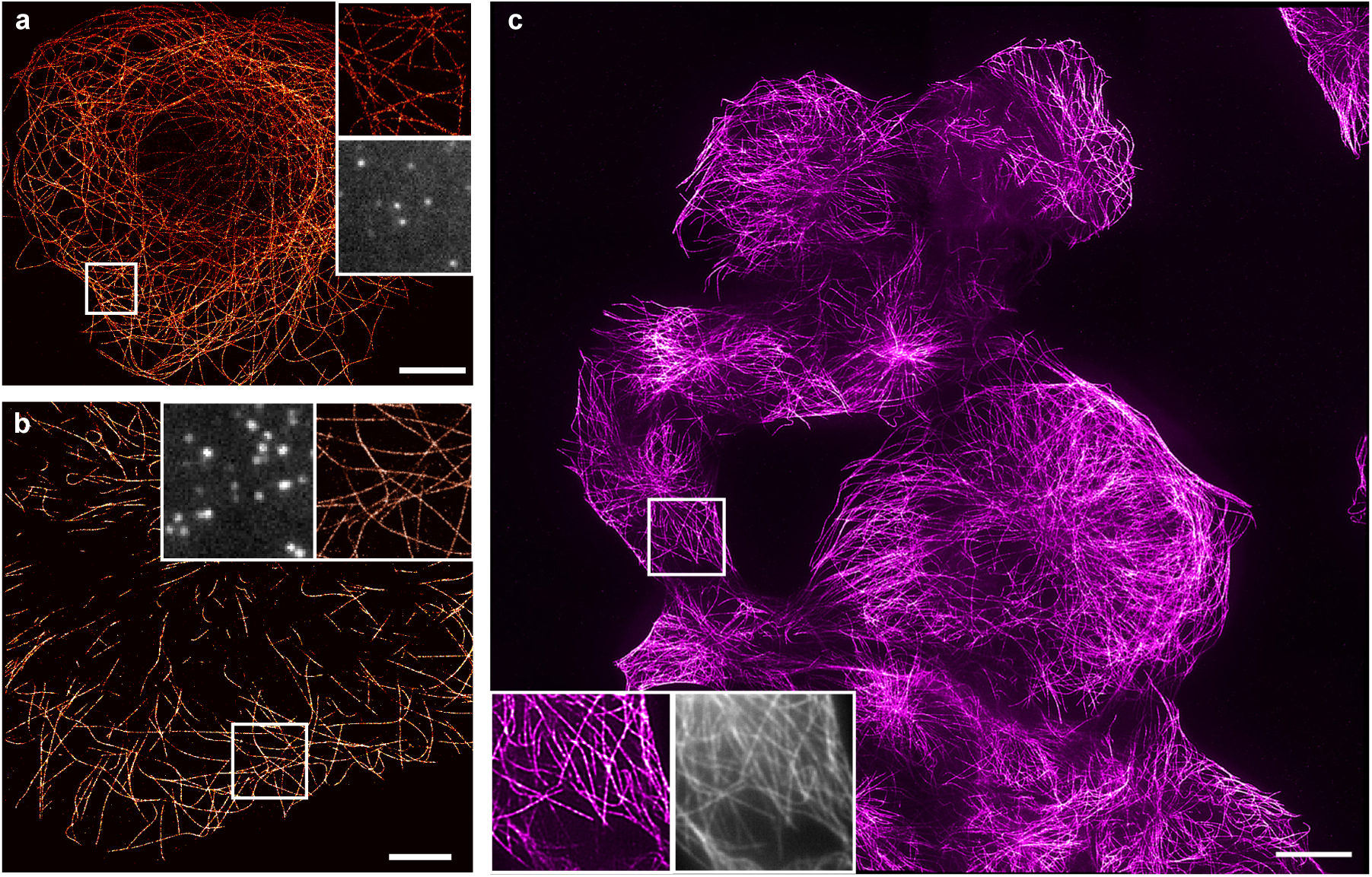
Super-resolution images of microtubules in fixed COS-7 cells. Each image includes a zoom-in area (colored) and a raw image frame of the same ROI (grayscale). **(a)** dSTORM with label of Alexa Fluor 647. 10 000 frames at 70 ms exposure time and 1.2 kW/cm^2^ laser intensity at 638 nm. Localization precision = 8.21 nm, scale bar = 10 *µ*m; **(b)** DNA-PAINT with label of P1 docking strand. 10 000 frames at 100 ms exposure time and 0.39 kW/cm^2^ laser intensity at 561 nm. Localization precision = 13.0 nm. Scale bar = 5 *µ*m; **(c)** Second order SOFI with label of 5*×*R1 DNA-PAINT docking strand, stitching of 4 by 4 tiles, each tile reconstructed with 500 frames at 10 ms exposure time and 0.59 kW/cm^2^ laser intensity at 568 nm. Scale bar = 20 *µ*m.

SOFI does not require localization of individual molecules and instead improves the resolution n times by n-th order cumulant analysis of fluorescence fluctuation.^6^ Using the blinking speed-optimized DNA-PAINT docking strand 5*×*R1 and a high imager strand concentration at 20 nM,^32^ we could reconstruct a second-order SOFI with 500 frames of dense and fast blinking data.^33^ By scanning through a large FOV consisting of a 4 by 4 grid using *µ*Manager’s multi-position acquisition, high-throughput super-resolution imaging was achieved after stitching the second-order SOFI images reconstructed from each grid, shown in Fig.4. The stitched FOV size is 0.23 mm *×* 0.23 mm and acquired within a total camera exposure time of 80 ms.

## 3 Discussion

Open Blink bridges the gap between state-of-the-art quantitative super-resolution imaging performance and broad accessibility. By combining experimentally validated optical performance, including homogeneous large-field illumination, sub–10 nm localization precision, and nanometric axial stability over multi-hour acquisitions, with modular and cost-effective hardware, we demonstrate that high-end SMLM capabilities are not inherently restricted to proprietary commercial platforms. Rather than prioritizing affordability at the expense of performance, Open Blink was designed to maintain the key optical requirements for quantitative single-molecule localization microscopy while reducing technical and financial barriers for adoption.

The compact quad-line laser combiner provides uniform high-power excitation for SMLM while enabling homogeneous excitation conditions compatible with spatially consistent localization performance. In addition, it can serve as an add-on excitation source to upgrade a conventional wide-field microscope for super-resolution microscopy, or to enable imaging methods that benefit from homogeneous high-power illumination, such as quantitative fluorescence imaging and single-particle tracking. The affordable diode lasers contribute largely to the overall cost reduction of Open Blink, which could be further lowered with emerging green semiconductor laser sources.

Previous implementations of MiCube in TIRF microscopy used single mode lasers. We showed the compatibility of TIRF illumination with multimode optical fibers in MiCube by reducing its width and using a shorter 150 mm lens, which simultaneously enlarges the field of view size. The smaller MiCube led to our design of an adaption plate for both the sample stage and an enclosure chamber without interfering with the optics around the body. This allows the straightforward adoption of environmental control for future live-cell imaging, for example with previously reported cost-effective open solutions.^34^

The complete integration of Open Blink in *µ*Manager is a main advantage for users who wish to avoid programming work for microscope control and image acquisition. The firmware and the *µ*Manager plugin for our laser control can be readily generalized to control any device from *µ*Manager that supports 0–5 V analog modulation or pulse-width modulation through a micro-controller. The plugin enabled metadata writing, addressing this limitation of previous micro-controller plugins available for *µ*Manager. Our infrared laser based *fg*Focus fills the missing gap of a *µ*Manager compatible focus-lock module. The *fg*Focus firmware and plugin can be easily transferred to other sample stages with an application programming interface (API) similar to that of the SmarAct stages. With the large library of hardware plugins existing for *µ*Manager, Open Blink can be deployed as a platform for highly automated smart microscopy.^35,36^ On-the-fly image analysis and microscopy control require informed image acquisition. We prepare Open Blink for such applications with metadata representing device status and acquisition parameters. For example, the laser power and TIRF stage position associated with each single molecule image can be used to actively modulate the blinking density of single fluorophores.

At substantially reduced cost around 70 000 euros, approximately one quarter of a commercial system for single-molecule localization microscopy, Open Blink demonstrates performance metrics comparable to high-end commercial systems in dSTORM, DNA-PAINT and SOFI imaging. We designed customized mechanical parts for simple in-house machining, whereas the electronics are ready-to-order with PCB files. The remaining high-cost components, such as the high-NA objective, sCMOS camera and the nanometer precision sample stage, can be replaced by lower cost alternatives for specific applications that tolerate the trade-off performance limitations.^13,37^

In summary, Open Blink demonstrates that high-performance quantitative single-molecule super-resolution microscopy can be made broadly accessible without compromising optical quality. By pairing validated optical performance with modular, affordable hardware and fully integrated open-source software, Open Blink lowers the barrier for laboratories to adopt state-of-the-art SMLM techniques. With comprehensive documentation and ready-to-use software, Open Blink not only expands access to super-resolution microscopy but also encourages innovation and development within the open hardware and software community.

## 4 Materials and Methods

### 4.1 Laser combiner construction

All lasers mentioned in Section 2.1 are mounted on a customize-cut aluminum heat sink, which adjusts all beams to the same height of 70 mm. Thermal paste is applied between the laser base plate and the heat sink to ensure efficient heat transfer. The lasers are positioned in parallel in the order of wavelength. Two 638 nm lasers are oriented in perpendicular to each other and are combined into a single beam by a polarizing beam splitter (PBS251, Thorlabs) mounted on kinematic mount (KM100PM/M, Thorlabs). Three longpass dichroic mirrors (DMLP425/DMLP550/DMLP605, Thorlabs) on kinematic mirror mounts (MDI, Radiant Dyes) sequentially combine beams of different wavelengths.

The combined beam is launched into an MMF (NA 0.22, square core profile of size 70 µm by 70 µm, customized, Ceram Optec) with a FC/PC connector (SM1FC, Thorlabs) via a 19 mm aspherical lens (AC127-019-A, Thorlabs) which is mounted in an adjustable lens tube (SM05V10, Thorlabs). Homogeneity via MMF agitation is facilitated with the use of a vibrational motor (5mm - 11mm Type, 304-111, precision microdrives) in proximity to the optical fiber within a 3D printed mount (customized, CAD available). The laser beam is coupled out by a FC/PC connector, after which an achromatic lens of 30 mm (AC254-030-A-ML, Thorlabs) collimates the beam before a laser clean-up filter (ZET 405/488/561/640xv2, Chroma) removing the autofluorescence from the MMF.

After the collimation lens, the beam size reducer consisting of two aligned achromatic lenses can be used for the small FOV configuration. The two lenses (AC254-050100-A-ML and ACN254-050-A, Thorlabs) mounted in lens tubes can be screwed onto two cage plates (CXY1A, Thorlabs) on the same cage assembly where the MMF adapter and the laser collimation lens are mounted on, such that beam size is halved in diameter and remains collimated and aligned along the optical axis. When larger FOV is needed, the two lenses can be taken down without changing the cage plates positions.

### 4.2 Microscope construction

The construction of Open Blink after the MMF exit is described as follows. In the excitation path, the collimated excitation beam is walked by two elliptical mirrors (BBE1-E02, Thorlabs), after which a 150 mm achromatic lens (AC254-150-A-ML, Thorlabs) focuses the beam onto the BFP of the objective. The precise positioning of the beam on the back focal plane of the objective is facilitated by a linear motorized stage (KMTS25E/M, Thorlabs) where the second elliptical mirror and the 150 mm focusing lens are mounted upon. Sample stage with nanometer precision (assembled with three identical linear stage, CLS5252, SmarAct) is mounted on a customized adaption plate (aluminum, customized, CAD available) on top of the modified MiCube (aluminum, customized, CAD available). An oil objective (UPLAPO60XOHR, Olympus) is screwed down at a fixed position. Fluorescence collected by the objective is decoupled by a quad-band dichroic mirror (zt405/488/561/640rpc, Chroma) mounted below (DFM1/M, Thorlabs) the objective and inside the miCube. A right angle mirror (KCB1E/M, Thorlabs) directed the fluorescence to be focused by the 180 mm tube lens (TTL180-A, Thorlabs) forming the intermediate image. At the focus of the tube lens, an adjustable slit (VA100/M, Thorlabs) is placed in order to restrict the image size for dual-color imaging. A quad-band filter before the tube lens (ZET405/488/561/640mv2, Chroma) filters out residual excitation light.

Further, the detection path features a dual-channel scheme. A 300 mm achromatic lens (G322-336322, Qioptiq) in 4f configuration with the tube lens creating the optical infinity space, allowing for insertion of a long pass dichroic mirror (ZT640rdc-UF3, Chroma) for color splitting, and of a square mirror (BBSQ1-E02, Thorlabs) for maximum collection of the fluorescence passing through the dichroic mirror. Both the dichroic mirror and the mirror are glued on a compact kinematic mount (KMS/M, Thorlabs) with silicone glue (Twinsil, Picodent). The two channels both include a 300 mm achromatic lenses (G322336322, Qioptiq) focusing the short and long wavelength images side by side onto the sCMOS camera (BSI Express, Teledyn Photometrics). In order to co-focus the two channels, the achromatic lens in the long wavelength channel is mounted onto a small dovetail stage (DT12/M, Thorlabs) allowing for fine adjustment along the optical axis. Proper bandpass filters can be inserted in either channel according to the fluorophore used.

The focus-lock module *fg*Focus utilizes an 830 nm infrared laser (CPS830S, Thorlabs) at TIRF configuration whose reflection on a linear sensor indicates the position of the sample plane. The infrared beam is first focused by a plano-convex lens (LA1908-B-ML, Thorlabs) and relayed by two mirrors (BBE1-E03, Thorlabs), the second one of which is translated by a 1D manual stage (DTS25/M, Thorlabs) for TIRF. The beam is then coupled into the microscope by a dichroic mirror (zt 775 sp-2p, AHF) overlapping the main excitation path after the TIRF focusing lens. The reflected infrared beam from the sample is decoupled by a 50/50 beam splitter (CCM1-BS014/M, Thorlabs) before being further filtered and focused onto the detection PCB. The reduced MiCube size means less space left in between the excitation TIRF focusing lens and the microscope body, posting challenges into inserting a dichroic mirror coupling the infrared beam into the microscope. We solved this by gluing the dichroic mirror with the silicone glue on a compact kinematic mirror mount (KMS/M, Thorlabs).

### 4.3 Sample preparation

#### 4.3.1 Cell culture and immunostaining

COS-7 cells (DSMZ GmbH) were cultured in Dulbecco’s modified Eagle medium (DMEM) in high glucose (Thermo Fisher) with addition of 10% fetal bovine serum (FBS; Gibco, Thermo Fisher), 1% L-glutamine (Gibco, Thermo Fisher), 1% sodium pyruvate (Gibco, Thermo Fisher), and 1% penicillin-streptomycin (Gibco, Thermo Fisher). Cells were seeded one day before immunostaining on coverslips (#1.5 type, Carl Roth) within 6-well plates filled with the culture medium at 37*^◦^*C and 5% CO_2_.

24 h after seeding, the cells were firstly treated for 90 s at room temperature in a pre-warmed extraction buffer which bases on the microtubule-stabilizing buffer Kapitein (80 mM PIPES, 7 mM MgCl_2_, 1 mM egtazic acid, 150 mM NaCl, 5 mM d-glucose) with addition of 0.3% (v/v) Triton X-100 and 0.25% (wt/vol) glutaraldehyde. After the removal of the extraction buffer, the cells were fixed by the fixation buffer containing 4% (wt/vol) paraformaldehyde in PBS for 10 min at room temperature, followed by three 5-min washes with PBS on an orbital shaker. To quench the autofluorescence, the fixed cells were treated for 7 min under 10 mM freshly dissolved sodium borohydride in PBS and immediately rinsed with PBS before two more 10-min washes. The fixed cells were then permeabilized with 0.25% (v/v) Triton X-100 in PBS for 7 min and then washed for three times with PBS. Afterwards, the cells were blocked for an hour at room temperature with blocking buffer containing 2% (wt/vol) bovine serum albumin, 10 mM glycine, and 50 mM NH_4_Cl.

Cells were firstly immunostained with primary antibody against *α*-tubuluin at 200 times dilution (T5168-.2ML, Merck). For dSTROM in Fig.4 (a), cells were then incubated with anti-mouse secondary antibodies conjugated with Alexa Fluor 647 (A-31571, Thermo Fisher); For DNA-PAINT in Fig.4 (b), cells were incubated with anti-mouse secondary antibody conjugated with P1 docking strand (sequence 5’-3’, TTATACATCTA, Massive Photonics). For DNA-PAINT-SOFI in Fig.4 (c), cells were incubated with anti-mouse secondary nanobody conjugated with 5*×*R1 docking strand(sequence 5’-3’: TCCTCCTCCTCCTCCTCCT). Site-specific secondary nanobody conjugation with the docking strands is described elsewhere.^33^ Both secondary nanobodies or antibodies were diluted 200 to 300 times with the blocking buffer and applied to the fixed cells on coverslips for one-hour incubation time at room temperature, followed by three 5-min washes with blocking buffer. Finally, samples were incubated for 15 min in post-fixation buffer (2% paraformaldehyde in PBS), and stored in PBS at 4*^◦^*C until imaging.

#### 4.3.2 Imaging buffer and imaging acquisition

Fixed cells were imaged at room temperature in imaging buffer. For dSTORM imaging, GLOXY buffer with the following components was made freshly on the day of imaging: 33 mM cysteamine, 500 *µ*g/mol glucose oxidase, 40 *µ*g/mL catalse, 10% (w/v) glucose in 50 mM Tris buffer. For DNA-PAINT imaging, buffer consists of 1nM P1 imager (sequence 5’-3’: AGATGTAT-Cy3B) in PBS with 500 mM NaCl. For DNA-PAINT-SOFI imaging, buffer consists of 20nM R1 imager (sequence 5’-3’: AGGAGGA-Atto655) and 5% ethlyne carbonate in PBS with 500 mM NaCl. Imaging acquisition is through *µ*Manager 2.0. For the large FOV DNA-PAINT-SOFI image, 4 by 4 tiles are acquired through multi-position acquisiton function with 10% pixel overlap.

### 4.4 Data analysis

SMLM reconstruction for DNA-PAINT and dSTORM data was done in Picasso software (v0.9.5).^31^ Localization precision is estimated by Picasso’s integrated NeNA algorithm.^38^ For dSTORM imaging, raw data frames were subject to Fast Temporal Median Filter before SMLM reconstruction.^39^ SOFI reconstruction was carried out with customized MATLAB script available in https://www.github.com/kgrussmayer/sofipackage for individual tile in the large FOV, afterwards SOFI images were stitched together by *Stitching* plugin on ImageJ software.

## Code, Data, and Materials Availability

A virtual CAD model of Open Blink, a detailed list of components used and their prices, design files of the customized parts related to Open Blink, as well as raw image data underlying Fig.4, are available in the online repository 4TU.ResearchData under the license CERN-OHL-S-2.0. An overview of the project is hosted at Grußmayer Lab GitHub page. *µ*Manager plugins with instructions are available at https://github.com/GrussmayerLab/LaserController and https://github.com/GrussmayerLab/fgFocus.

## Acknowledgments

The authors thank N. van Vliet for cell culture work, and S.-T. Hung for help with microscope alignment. The authors acknowledge D. de Roos and the DEMO workshop at TU Delft for their help in mechanical machining. The authors are grateful to C. Niederauer from Kristina Ganzinger lab, and all members of the Grussmayer lab for advice and discussions. S. El-Gebali gave the authors insightful guidance for open science practices. This work was supported by the TU Delft Department of Bionanoscience and the TU Delft Bioengineering Institute, and by the TU Delft Open Research Hardware Stimulation Fund to R.H. and to J.K.. K.G. was supported by an ERC grant (QScope, 101165129). Views and opinions expressed are however those of the author(s) only and do not necessarily reflect those of the European Union or the European Research Council Executive Agency. Neither the European Union nor the granting authority can be held responsible for them.

## Author Contributions

K.G. conceived the idea, initiated and supervised the project. K.G. and R.H. designed the experiments. R.H. assembled the microscope, performed experiments and analyzed the data. M.E. assisted R.H. in alignment and documentation. R.H. and J.K. designed the control electronics. J.K. implemented the software plugins. A.M. and J.E. initialized the design of data acquisition for the focus-lock module. R.H. and K.G. wrote the manuscript with input from all the other authors.

