## Supplementary Information for "Open Blink: Low-cost TIRF microscopy for super-resolution imaging via *µ*Manager"

[a]Department of Bionanoscience, Delft University of Technology, Van der Massweg 9, 2629 HZ Delft, the Netherlands
[b]Kavli Institute of Nanoscience Delft, Lorentzweg 1, 2628 CJ Delft, the Netherlands
[c]Haute école du paysage, d'ingénierie et d'architecture de Genève (HEPIA), HES-SO - Haute Ecole Spécialisée de Suisse Occidentale, Rue de la prairie 4, 1202 Geneva, Switzerland

### Supplementary information 1

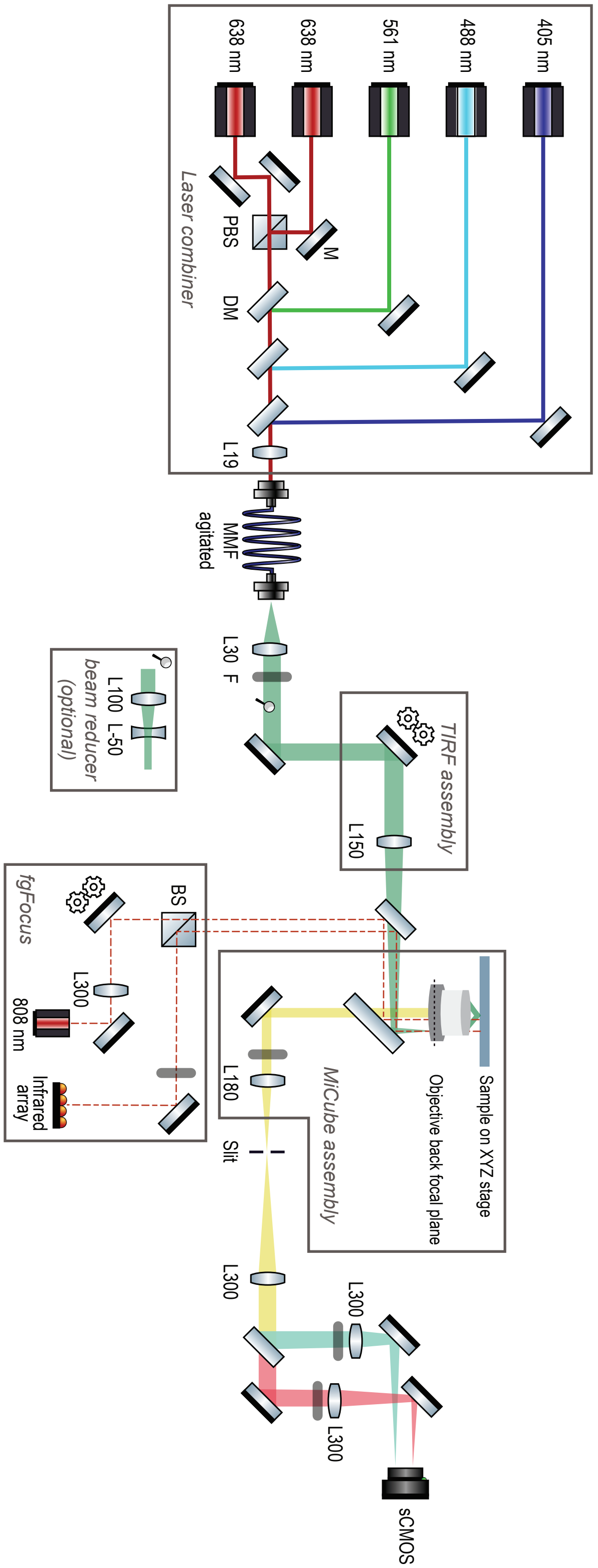


**Fig S1** Optical path of Open Blink. M: mirror. BS: beam splitter. PBS: polarized beam splitter. DM: dichroic mirror. F: filter. MMF: multimode fiber.

### Supplementary informantion 2

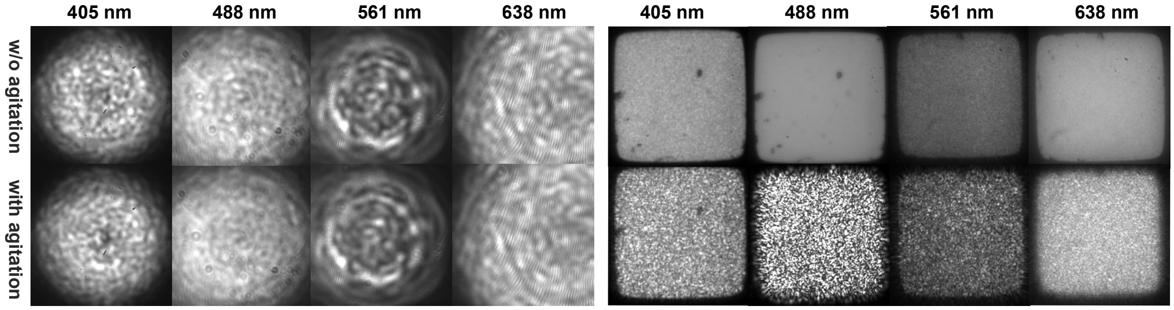


**Fig S2** Two alternative MMFs respond differently to the agitation. Images taken at the MMFs exits with 30 or 50 ms exposure time show laser intensity profiles at different wavelengths in the presence and absence of agitation. Left: circular-core MMF (M42L05, Thorlabs) with diameter of 50 $\mu$m; right: square-core MMF (M103L05, Thorlabs) sized 150 $\mu$m by 150 $\mu$m.

**Supplementary information 3**

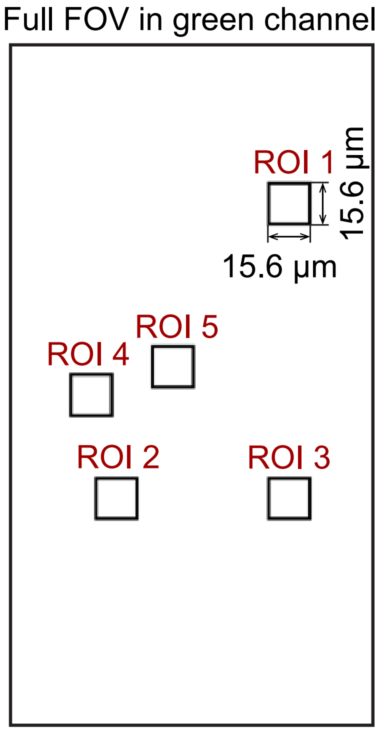


**Fig S3** Illustrated positions of region of interests (ROIs) used in DNA-PAINT imaging for evaluating sample plane laser homogeneity.

**Supplementary information 4**

The viability of TIRF condition is calculated according to the method reported in Kwakwa et al.[1] In TIRF, the excitation beam is focused within the TIRF annulus at the back focal plane of the objective whose width $S_a$ is defined by the objective parameters, i.e. the effective focal length EFL and numerical aperture (NA), in addition to the refractive index of the sample ($n_s$, 1.335 for PBS): $S_a$ = EFL×(NA-$n_s$). Our 60× oil objective with EFL of 3 mm and NA of 1.5 leads to $S_a$ = 495 $\mu$m of critical beam size for TIRF. The theoretical focused beam size $S_b$ defined by the MMF core diameter ($S_c$ = 70 $\mu$m), and the combination of the MMF collimation lens ($F_{cl}$ = 30 mm) and focusing lens ($F_k$ = 150 mm) as in $S_b$ = $S_c \times F_k/F_{cl}$ which equals 350 $\mu$m, smaller than $S_a$ thus allowing the realization with TIRF in our design.

### Supplementary 5

**Table S1** Sample plane excitation intensity at the two different FOV configurations. The maximum laser power of each wavelength is measured directly above the objective under epi-illumination. The FOV size is measured by the camera image with a dense fluorescent beads sample under epi-illumination. Note that the measured FOV sizes are slightly smaller than the theoretical size determined by the excitation optics ($264 \times 264\ \mu m^2$ for the large FOV configuration and $132 \times 132\ \mu m^2$ for the reduced FOV configuration), due to measurement error and the non-collimated component in the MMF output.

| Maximum power density at sample plane ($kW/cm^2$) | 405 nm | 488 nm | 561 nm | 638 nm |
|---|---|---|---|---|
| FOV = $257 \times 257\ \mu m^2$ | 0.67 | 0.7 | 0.67 | 1.09 |
| FOV = $105 \times 105\ \mu m^2$ | 4.71 | 4.83 | 4.67 | 6.94 |

*References*


1 K. Kwakwa, A. Savell, T. Davies, *et al.*, “easySTORM: a robust, lower-cost approach to localisation and TIRF microscopy,” *Journal of Biophotonics* **9**, 948–957 (2016).
